# Optogenetic control of the Bicoid morphogen reveals fast and slow modes of gap gene regulation

**DOI:** 10.1101/2021.10.13.464280

**Authors:** Anand P. Singh, Ping Wu, Sergey Ryabichko, João Raimundo, Michael Swan, Eric Wieschaus, Thomas Gregor, Jared E. Toettcher

## Abstract

Developmental patterning networks are regulated by multiple inputs and feedback connections that rapidly reshape gene expression, limiting the information that can be gained solely from slow genetic perturbations. Here we show that fast optogenetic stimuli, real-time transcriptional reporters, and a simplified genetic background can be combined to reveal quantitative regulatory dynamics from a complex genetic network *in vivo*. We engineer light-controlled variants of the Bicoid transcription factor and study their effects on downstream gap genes in embryos. Our results recapitulate known relationships, including rapid Bicoid-dependent expression of *giant* and *hunchback* and delayed repression of *Krüppel*. In contrast, we find that the posterior pattern of *knirps* exhibits a quick but inverted response to Bicoid perturbation, suggesting a previously unreported role for Bicoid in suppressing *knirps* expression. Acute modulation of transcription factor concentration while simultaneously recording output gene activity represents a powerful approach for studying how gene circuit elements are coupled to cell identification and complex body pattern formation *in vivo*.

## Main Text

Gene networks play a crucial role in developmental patterning, transforming rudimentary positional cues into a multitude of sharply defined domains of gene expression. Such networks are typically characterized by redundant inputs to ensure that gene expression is initialized appropriately, and feedback connections between genes in the network to ensure a consistent patterning response. Information from these inputs is integrated at enhancers that bind multiple transcription factors and control gene expression through transient interactions with promoters and longer-term alterations of chromatin structure and accessibility. Understanding how networks function requires knowing the time scales over which individual components operate.

The gap gene network of the early *Drosophila* embryo is a canonical example of such a sophisticated pattern-forming system. In this network, the expression of four core transcription factors – the gap genes *giant (gt), hunchback (hb), Krüppel (Kr)*, and *knirps (kni)* – is initiated by three partially-redundant sources of positional information that are maternally deposited in the egg. These maternally supplied inputs include an anterior-to-posterior gradient of the Bicoid (Bcd) transcription factor, a posterior-to-anterior gradient of the Nanos RNA binding protein, and Torso receptor tyrosine kinase signaling at the anterior and posterior poles. In addition to responding to maternal inputs, the four gap genes further regulate themselves and each other to generate bands of gene expression that are essential for specifying the body plan (**Figure 1a-b**)^1^.

**Figure 1.**
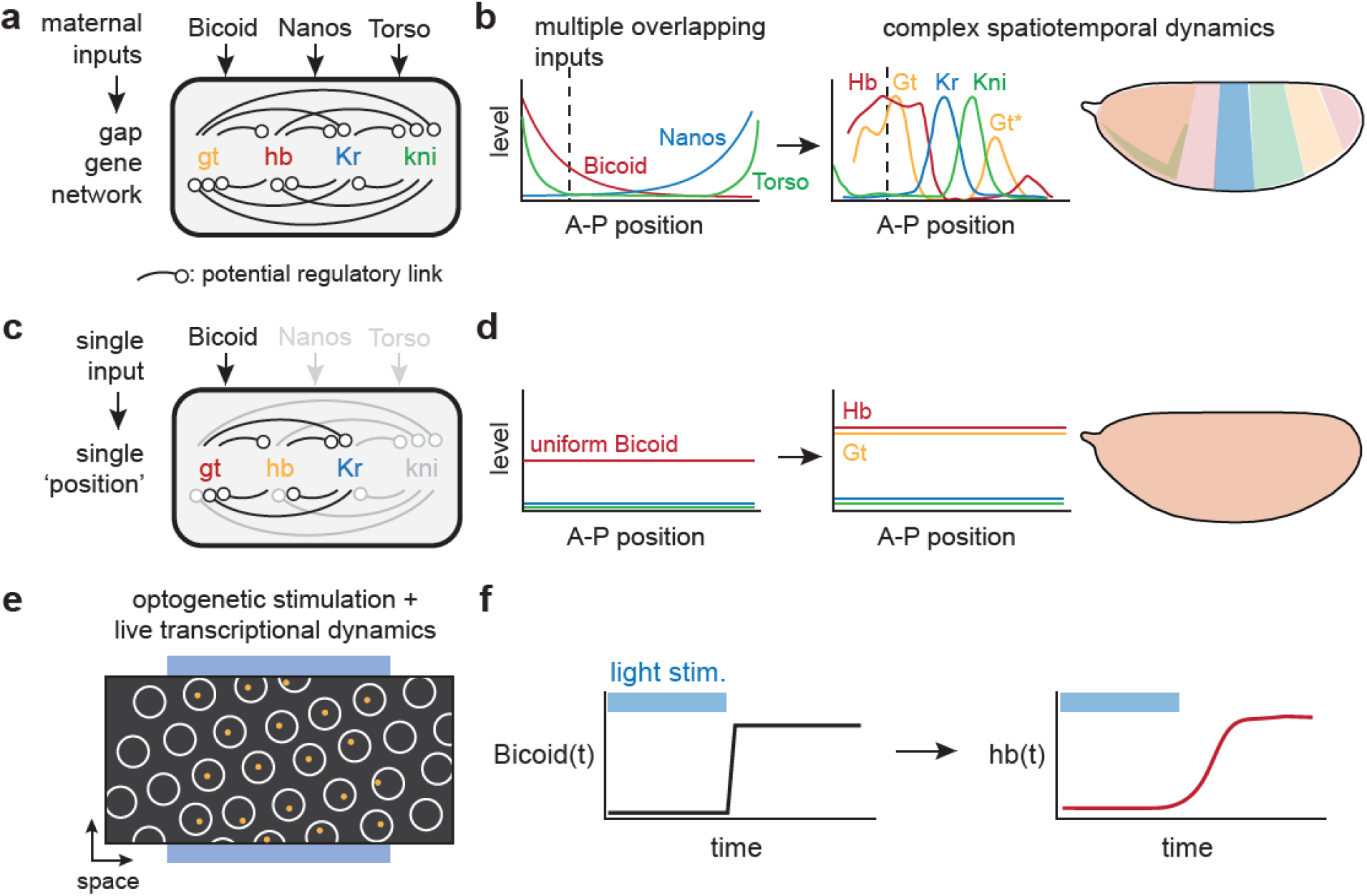
Studying Bicoid-dependent gap gene responses using a stimulus-response approach in single-input embryos. (**a-b**) The endogenous gap gene network depends on three maternally-supplied inputs (Bicoid, Nanos, and Torso) and many potential feedback and crosstalk links (in **a**) to generate bands of gap gene expression across the embryo (in **b**). A detailed understanding of this network is made challenging by the presence of multiple redundant inputs and complex dynamics as spatial patterns shift over time. (**c-d**) To study the effects of the Bicoid transcription factor on gap gene expression, we set out to construct a reduced-complexity network where Bicoid is the sole maternally supplied input to the network (in **a**) and spatial patterning is eliminated (in **b**), eliminating both redundancy and spatiotemporal dynamics. (**e-f**) To define the strength, duration, and dynamics of Bicoid-dependent gap gene responses, we acutely perturb nuclear Bicoid levels using an optogenetic technique and monitor resulting gene expression in individual nuclei using live transcription reporters.

A useful first step in disentangling such networks has been to characterize transcriptional responses under conditions where input information has been reduced to single components or flattened so that all cells in the embryo see the same input values (**Figure 1c-d**). For example, when all anterior-posterior (A-P) patterning inputs except Bicoid are eliminated, the pattern is reduced and shifted relative to wildtype, but the fundamental sequence of all four gap genes is maintained^2^. Similarly, flattening spatial patterns generates embryos that reflect a single A-P position along the wild-type gradient without the complexity of network components diffusing across shifting gene expression boundaries^3,4^. While such simplified systems can provide useful insights about Bicoid-dependent features of the network, they do not distinguish between direct and indirect effects, or long- and short-term mechanisms.

Real-time measurement of responses to acute perturbations provides a useful approach to characterize a network’s dynamic features^5,6^. Differences in response kinetics can also distinguish direct interactions (e.g., where a transcription factor directly regulates its target’s expression) from indirect links (where an intermediate gene product must first be synthesized). Although such rapid stimulus-response experiments have been traditionally difficult to perform *in vivo*, the recent advent of optogenetic perturbations and live biosensors of gene expression offer the possibility to dissect gene expression networks with unprecedented resolution (**Figure 1e-f**)^6-10^.

Here, we describe a strategy for dissecting Bcd-dependent gap gene responses using fast stimulus-response measurements in simplified embryos that lack redundant inputs and spatial patterns. We generated a series of light-sensitive Bcd variants whose nuclear-cytosolic localization can be shifted in less than a minute using blue light. We introduced these variants in embryos that lack all other sources of A-P asymmetry, eliminating the protein gradients and shifting spatial distributions that typically complicate the study of patterning gene networks. When constructed using optogenetic Bcd variants with different activity levels, these synthetic spatially-homogeneous embryos mimic either anterior, central, or posterior embryonic positions, offering a toolbox for studying the real-time transcriptional responses to acute perturbation of a developmental patterning cue. Combining acute optogenetic Bcd perturbation with live-embryo biosensors of gap gene expression reveals both rapid and delayed modes of Bcd-dependent regulation. Anteriorly expressed gap genes *gt* and *hb* respond within minutes to changes in Bcd concentration, consistent with a direct role for Bcd in their transcriptional activation. In contrast, the medial gap gene *Kr* exhibits a delayed and inverted response, indicative of indirect Bcd-induced repression through an intermediate node. Finally, we report that the posteriorly expressed gap gene *kni* is rapidly transcribed upon acute loss of nuclear Bcd, an unexpected response suggesting that Bcd acts to repress *kni* expression without requiring new gene synthesis. Our approach, combining rapid nuclear-cytosolic shuttling of a transcription factor with real-time transcription measurements in a developmental gene regulatory network, offers the possibility to dissect regulatory links with unprecedented precision.

## Results

### An activity series of optogenetic Bicoid variants with rapid stimulus-response kinetics

We engineered optogenetic variants of Bcd to serve as rapidly-switchable inputs to the gap gene network. We fused Bcd to LEXY, an optogenetic tool based on the AsLOV2 protein domain whose nuclear export is reversibly triggered by blue light, enabling rapid and titratable optogenetic control over transcription factor activity^11-15^. Blue light illumination uncages a buried nuclear export sequence (NES) in LEXY’s C-terminal Jα helix; in the dark, NES activity is lost and Bcd’s nuclear localization signal (NLS) returns the fusion protein into the nucleus (**Figure 2a**). LEXY-based translocation typically produces a 5-fold change in nuclear protein concentration^11,13^ whereas the natural Bcd gradient is thought to affect gene expression over a larger range, suggesting that Bcd variants at various expression or activity levels may be required to probe gap gene responses at different embryonic positions. We thus tested a series of Bcd-LEXY variants that were either left untagged or N-terminally fused to different fluorescent proteins, a modification previously observed to generate distinct Bcd activity levels (see **Methods**)^16^.

**Figure 2.**
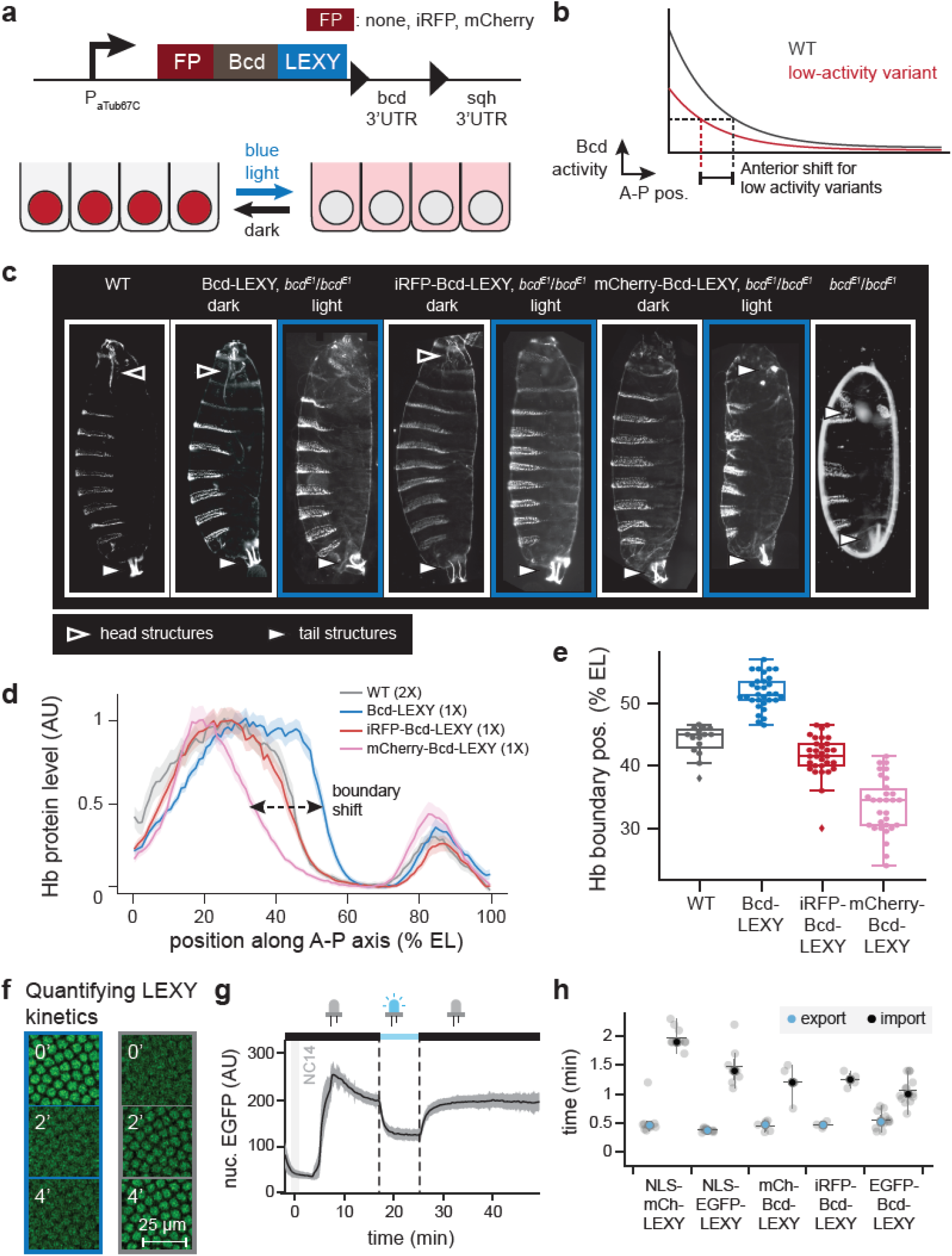
A series of optogenetic Bicoid variants with variable activity and rapid kinetics. (**a**) Bicoid was fused to various fluorescent proteins at its N terminus and to the LEXY optogenetic system at its C terminus. 450 nm light illumination exposes LEXY’s nuclear export sequence (NES), leading to an expected decrease in Bicoid transcriptional activity. (**b**) Bicoid variants harboring weaker activity are expected to exhibit loss of anterior structures and an anterior shift of gene expression patterns. (**c**) Larval cuticles for Bcd-LEXY variants in dark and light conditions. Anterior head and posterior tail structures are indicated with the outlined and shaded arrows. Illuminated embryos exhibit loss of anterior structures or duplication of posterior structures, indicating progressive loss of Bicoid activity. (**d-e**) Immunofluorescence for Hunchback (Hb) protein for three Bcd-LEXY variants, compared to WT embryos. Hb levels are quantified as a function of position and genetic background in **d**, with the boundary of anterior Hb expression quantified for individual embryos in **e**. Bcd-LEXY exhibits high activity, whereas iRFP-Bcd-LEXY and mCherry-Bcd-LEXY exhibit progressively weaker activity as determined by the boundary of anterior Hb expression. (**f-g**) EGFP-Bcd-LEXY time course during cycle of optogenetic activation and deactivation. Representative images are shown in **f**; quantification during a nuclear cycle 14 (NC14) time course of light and dark exposure shown in **g**. (**h**) Quantification of import and export kinetics for five LEXY constructs is shown in **h**; similar kinetics of translocation are observed for fluorescent Bicoid variants and non-Bicoid-containing LEXY constructs.

To assess the function of each Bcd-LEXY variant, we generated embryos harboring a single variant as the sole Bcd source and assessed its function in the light and dark. Bcd is normally expressed in an anterior-to-posterior gradient, so conditions in which Bicoid activity is reduced should lead to loss of anterior structures and/or an anterior shift of gap gene expression patterns (**Figure 2b**). Bcd-LEXY and iRFP-Bcd-LEXY embryos exhibited body segmentation and cephalic furrow position consistent with high Bcd activity in the dark, and embryos harboring a single copy of either allele hatched at rates of 70% and 42%, respectively (**Figure 2c**; **Table S1**). Blue light led to an apparent reduction in Bcd activity, characterized by the loss of mouth parts and thoracic segments at the anterior and a loss of embryo viability. mCherry-Bcd-LEXY carrying embryos displayed weaker overall Bcd activity, as these embryos failed to form anterior structures in the dark and phenocopied *bcd*^*E1*^ loss-of-function embryos under illumination (**Figure 2c**, right panels).

We also measured expression of the canonical Bcd target gene Hunchback (Hb) in embryos harboring each of the three Bcd-LEXY variants (**Figure 2d-e**). Compared to wild-type embryos, the position of the Hb boundary is shifted towards the posterior in Bcd-LEXY embryos and progressively shifts toward the anterior in iRFP-Bcd-LEXY and mCherry-Bcd-LEXY embryos, confirming that these three variants form an activity series: Bcd-LEXY > iRFP-Bcd-LEXY > mCherry-Bcd-LEXY. Immunostaining for Bcd revealed that the activity differences between our Bcd-LEXY variants are partially explained by differences in expression level between all three variants (**Figure S1a**). Nevertheless, all three variants were expressed at higher levels than wild-type Bcd, suggesting that fusion to LEXY and/or a fluorescent protein also partially interferes with Bcd function^16,17^.

We quantitatively characterized nuclear import and export dynamics for each Bcd-LEXY variant as well as two LEXY-tagged fluorescent proteins that lacked any transcription factor fusions (NLS-mCherry-LEXY and NLS-EGFP-LEXY) (**Figure 2f-h**; see **Methods** for imaging details). Switching 450 nm light on or off led to a rapid redistribution of each Bcd-LEXY variant in nuclear cycle 14 (NC14) embryos (e.g., EGFP-Bcd-LEXY in **Figure 2g**). Comparable dynamic responses were observed across all variants, with light-induced nuclear export in 30 sec and darkness-induced import in 1-2 min (**Figure 2h**). Illumination also produced nuclear export of similar magnitude and spatial precision, with a 4-fold change in nuclear concentration between dark and light conditions and a spatial precision of ∼10-12 μm (1-2 cells) (**Figure S1b-d**). These data establish the LEXY system as a tool for rapid modulation of nuclear transcription factor concentration during pre-gastrulation *Drosophila* embryogenesis.

### A reduced-complexity embryo for dissecting Bcd regulation of gap gene expression

Our goal is to use the fly embryo as a laboratory to measure stimulus-response functions for Bicoid’s regulation of gap gene expression. We thus sought to simplify the experimental system, eliminating redundant inputs to the gap gene network as well as the complex spatial patterns found in wild-type embryos. Recent studies^2,3,18^ established genetic strategies for producing embryos that lack all known sources of anterior-to-posterior variation, and we used these triple-mutant *bcd*^*E1*^ *nos*^*BN*^ *tsl*^*4*^ / *bcd*^*E1*^ *nos*^*l7*^ *tsl*^*4*^ (henceforth referred to as *bnt*) embryos as a starting point for introducing light-controlled Bcd-LEXY variants (see **Methods** for detailed information on fly stock and genetics). We also reintroduced uniform levels of a weak Nanos variant (*nos* TCEIIUC:AG)^19,20^ to suppress maternal Hb protein expression; the resulting embryos, termed *nos-tub bnt*, are devoid of all three A-P patterning cues and produce a posterior-like gene expression state throughout the embryo (**Figure S2a**). Onto this background we expressed a single Bcd-LEXY variant at a uniform level across the embryo to serve as the sole patterning input to the gap gene network^3^. Bcd-LEXY *nos-tub bnt* embryos can thus be thought of as representing a single embryonic “position” set by Bcd which can subsequently be perturbed using light (see e.g. **Figure 3a** for *Kr* in uniform mCherry-Bcd-LEXY embryos).

**Figure 3.**
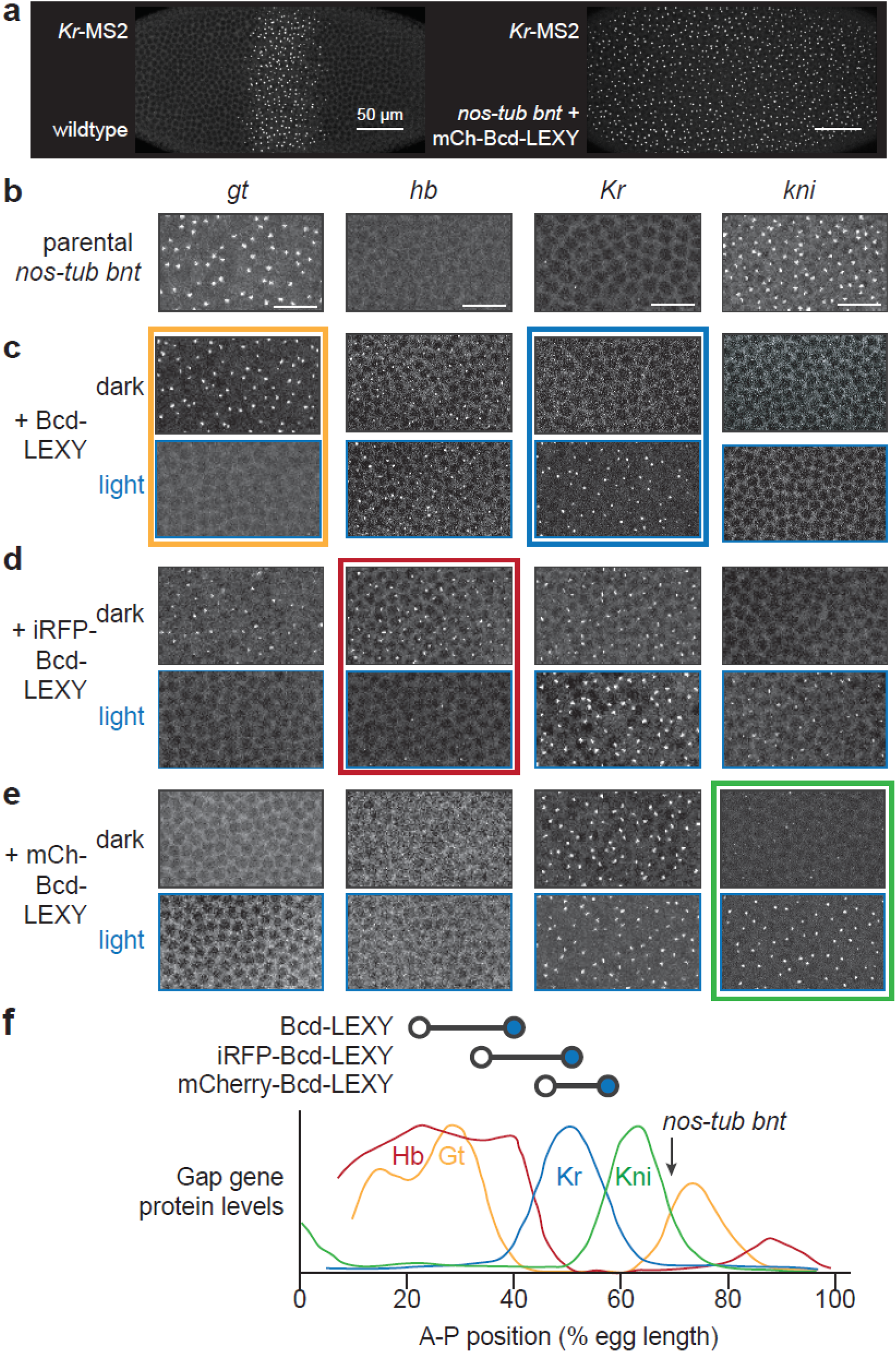
Spatially-uniform, single-input embryos to enable optogenetic interrogation of specific gap genes. (**a**) Nuclear cycle 14 embryos imaged using a *Kr* MS2 reporter. Left: embryo exhibiting wild-type A-P patterning. Right: *nos-tub bnt* embryo harboring a single copy of uniformly-expressed mCherry-Bcd-LEXY. The *nos-tub bnt* background eliminates all maternally supplied A-P patterns, so a uniformly expressed Bcd-LEXY variant produces a single approximate A-P position per embryo. (**b-e**) Regions of embryos showing MS2 reporter transcription for all four gap genes in (**b**) *nos-tub bnt*, (**c**) *nos-tub bnt* + Bcd-LEXY, (**d**) *nos-tub bnt* + iRFP-Bcd-LEXY, and (**e**) *nos-tub bnt* + mCherry-Bcd-LEXY embryos. For optogenetic illumination experiments, embryos were bathed in 450 nm light for 1 h. (**f**) Mapping approximate embryonic positions represented by dark and light conditions in each genetic background. Bottom: diagram from Ref. 2 quantifying gap gene expression as a function of A-P position, with posterior = 100% EL. Top: the approximate position based on gap gene expression for each optogenetic variant under in illuminated (open circle) and dark conditions (blue circle).

To define the A-P position represented by each uniformly expressed Bicoid-LEXY variant, we measured the transcriptional activity for four gap genes (*gt, hb, Kr* and *kni*) using the MS2/MCP system (see **Methods**; **Figure 3b-e**; **Figure S2b-c; Video S1-S4**). We found that our baseline *nos-tub bnt* embryos lacking all A-P patterning transcribed high levels of *kni* and *gt*, but low levels of *Kr* and no detectable *hb* (**Figure 3b** and arrow in **3f**). This pattern was altered dramatically in the presence of uniformly expressed Bcd-LEXY, which drove an anterior-like of *gt* and *hb* transcription in the dark, shifting to a mid-embryo-like state of *hb* and *Kr* transcription in the light (**Figure 3c**). Uniform iRFP-Bcd-LEXY embryos transcribed *gt, hb*, and *Kr* in the dark, shifting to *Kr* and *kni* expression in the light (**Figure 3d**). Finally, mCherry-Bcd-LEXY embryos shifted between weak and strong *kni* transcription depending on illumination conditions, with high *Kr* transcription in both cases (**Figure 3e**).

Comparing the combinations of gap genes expressed in each background to a wild-type embryo suggests a mapping between each Bcd-LEXY variant and embryonic position (**Figure 3f**). Illumination shifts Bcd-LEXY embryos from high *hb* and *gt* expression to *hb* alone, which normally occurs between ∼20-40% egg length (EL); similarly, iRFP-Bcd-LEXY embryos shift from ∼35-50% EL, and mCherry-Bcd-LEXY embryos from ∼45-60% EL (**Figure 3f**, top). Notably, parental *nos-tub bnt* embryos express high levels of *kni* and *gt*, consistent with a position of ∼70% EL, because their reduced expression levels of maternal Hb allows that expression but does not promote expression of more posterior targets in the absence of Torso/Erk signaling (**Figure 3f**, arrow). Importantly, these results define the optogenetic Bcd variants that can be used to switch each of the four core gap genes between high and low expression levels: *gt* (Bcd-LEXY), *hb* (iRFP-Bcd-LEXY), *Kr* (Bcd-LEXY) and *kni* (mCherry-Bcd-LEXY) (**Figure 3b-e**; colored boxes), which we used to interrogate the transcriptional dynamics of each gap gene in the following live-imaging experiments.

### Anterior patterns of *hb* and *gt* respond rapidly to changes in nuclear Bcd concentrations

How do gap genes respond to acute changes in nuclear Bcd concentration? To answer this question, we set out to combine optogenetic Bcd-LEXY control with live imaging of individual gap genes using the MS2/MCP system. We constructed a confocal microscope that combines a tunable 2-photon (2P) laser for GFP/mCherry imaging with a digital micromirror device and 450 nm LED for optogenetic stimulation (see **Methods**; **Figure 4a**; **Figure S3a**). 2P excitation is ideal because it can be used for simultaneous GFP and mCherry imaging without triggering AsLOV2 excitation (**Figure S3b**; **Video S5-6**), enabling high-resolution volumetric imaging without undesirable photoactivation of our optogenetic system^21,22^.

**Figure 4.**
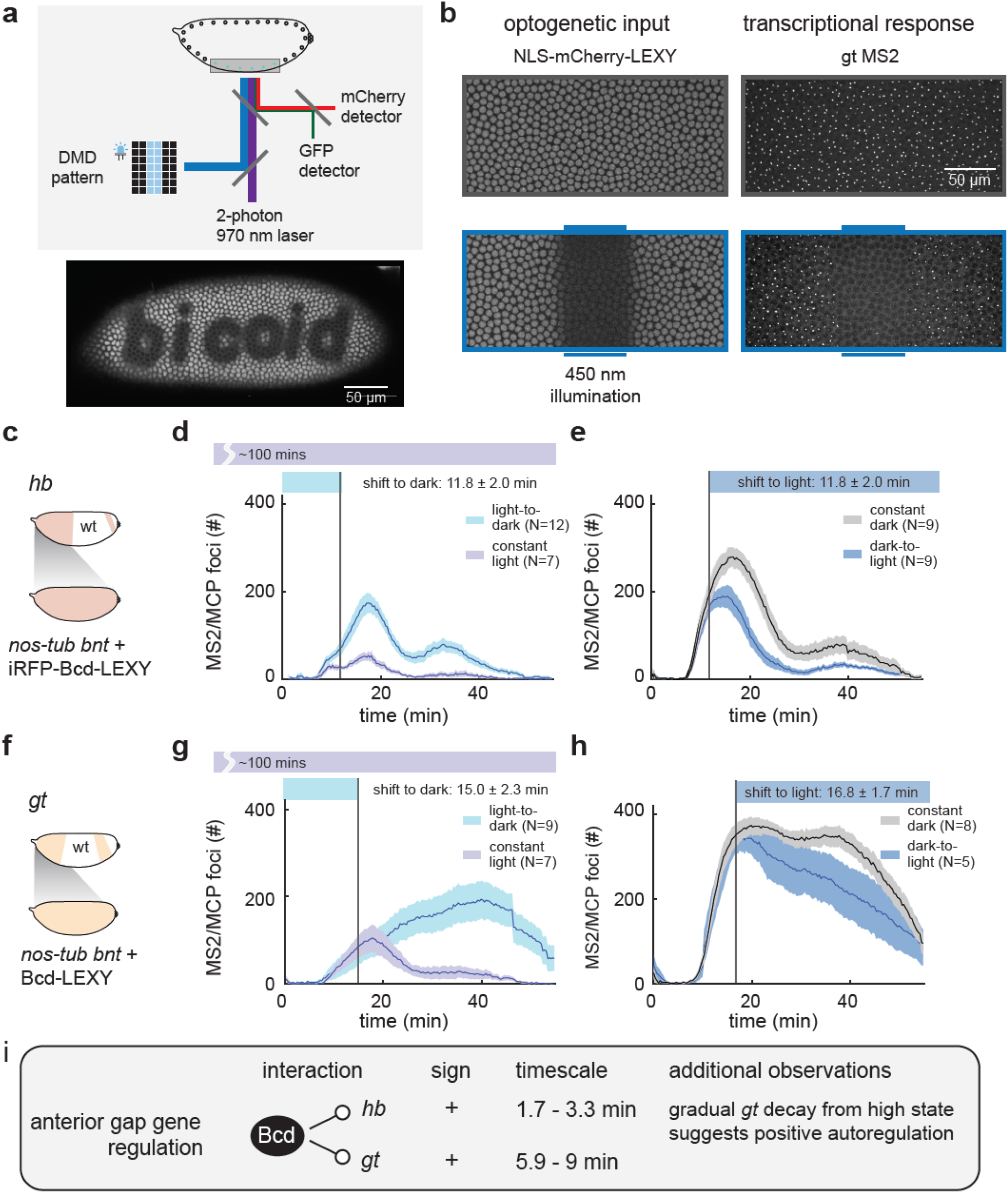
Optogenetic stimulation and live transcription measurement for anterior expression patterns of *gt* and *hb*. (**a**) Schematic of optogenetic activation and two-photon imaging setup. 450 nm LED light is patterned using a digital micromirror device (DMD) to deliver optogenetic stimuli. Two-photon imaging at 970 nm excites GFP and mCherry without crosstalk to the LEXY optogenetic system. (**b**) Example of light stimulation and two-color imaging of NLS-mCherry-LEXY and MCP foci for a *gt* MS2 transcriptional reporter. Images show ventral regions of representative embryos in the absence or presence of a 450 nm light input delivered in a stripe in the middle of the embryo. (**c-e**) Optogenetic interrogation of Bcd-induced anterior *hb* expression dynamics. Uniformly expressed iRFP-Bcd-LEXY embryos were imaged for *hb* MS2 reporter expression (schematic in **c**) upon an acute shift from light to dark (in **d**) and dark to light (in **e**); constant-light and constant-dark stimuli were sued as controls. (**f-h**) Optogenetic interrogation of Bcd-induced anterior *gt* expression dynamics. Uniformly expressed Bcd-LEXY embryos were imaged for *gt* MS2 reporter expression (schematic in **f**) upon an acute shift from light to dark (in **g**) and dark to light (in **h**); constant-light and constant-dark stimuli were used as controls. (**i**) Summary of stimulus-response results for *gt* and *hb*. Rapid light-triggered changes in both *gt* and *hb* transcription are consistent with direct activation by Bcd. Subsequent gradual changes in *gt* expression suggest the presence of positive autoregulation. For **d, e, g**, and **h**, shaded regions show standard error of the mean, and the number of embryos tested is indicated on each plot.

We engineered embryos that maternally express a desired uniform Bicoid-LEXY variant as well as two additional constructs: an MCP-mNeonGreen protein for live transcript visualization^23-25^ and an NLS-mCherry-LEXY indicator to define the current activity state of our optogenetic system (**Figure S3c-d**; see **Methods**). By crossing females of this genotype with males harboring a desired MS2-tagged gap gene reporter, we can thus deliver optogenetic stimuli while imaging both LEXY nuclear translocation and transcriptional responses in individual nuclei over time in live embryos (see **Figure 4b** using the *gt*-MS2 reporter).

We first performed stimulus-response measurements for Bcd regulation of anterior *hb* expression using our medium-activity iRFP-Bcd-LEXY variant (**Figure 4c-d**). We measured *hb* transcription in response to an acute increase in Bcd activity by shifting from blue light to dark conditions during early NC14; continuously illuminated embryos were used as a control (**Figure 4d**; **Figure S4a**). We found that *hb* expression rose rapidly after a light-to-dark shift; quantifying this response time revealed a shift within 1.7 ± 0.9 min after light perturbation (mean ± SEM; see **Table S2**). Conversely, acute removal of Bcd by switching from dark to light conditions caused *hb* expression to fall relative to dark-incubated controls within 3.3 ± 1.1 min (**Figure 4e**). These data indicate that gap gene transcription can respond extremely rapidly to acute increases or decreases in nuclear Bcd concentration and are consistent with the standard model of Bcd acting as a direct transcriptional activator of anterior *hb* expression.

How do the anterior expression dynamics of *gt* compare to those of *hb*? We examined a *gt* MS2 reporter^26^ under similar light-to-dark and dark-to-light illumination shifts in NC14 (**Figure 4f-h**; **Figure S4b**). As in the case of *hb*, we found that optogenetic Bcd perturbations were rapidly transmitted to *gt* expression, with response times of 5.9 ± 1.6 min and 9 ± 2.7 min depending on the illumination sequence, indicative of direct transcriptional activation (**Figure 4g-h; Table S2**). However, unlike the case for *hb, gt* transcription continued changing gradually, only approaching a final expression state over ∼30 min. A rapid initial response followed by delayed progression towards a final state is suggestive of positive autoregulation^27^, for example if *gt* transcription were positively influenced by the past history of gap gene (*gt* or *hb*) expression. Indeed, positive feedback on *gt* transcription by Gt protein was recently reported^26^. In summary, live-embryo stimulus-response measurements identify *hb* and *gt* as direct Bcd transcriptional targets and suggest *gt* as a potential target of positive autoregulation (**Figure 4i**).

### Optogenetic Bcd stimuli produce delayed and inverted *Kr* transcriptional responses

To explore how dynamic changes in Bcd concentration alter the expression of gap genes in the middle of the embryo, we next turned to the gap gene *Kr* (**Figure 5a**). *Kr* expression is known to be regulated by multiple transcription factors, including Bcd^28-30^, Stat92E^31^, Zelda (Zld)^32^, and Hunchback (Hb)^30,33^; this complex regulation is thought to ensure that *Kr* is expressed in a narrow central band, with low expression at both anterior and posterior embryonic positions.

**Figure 5.**
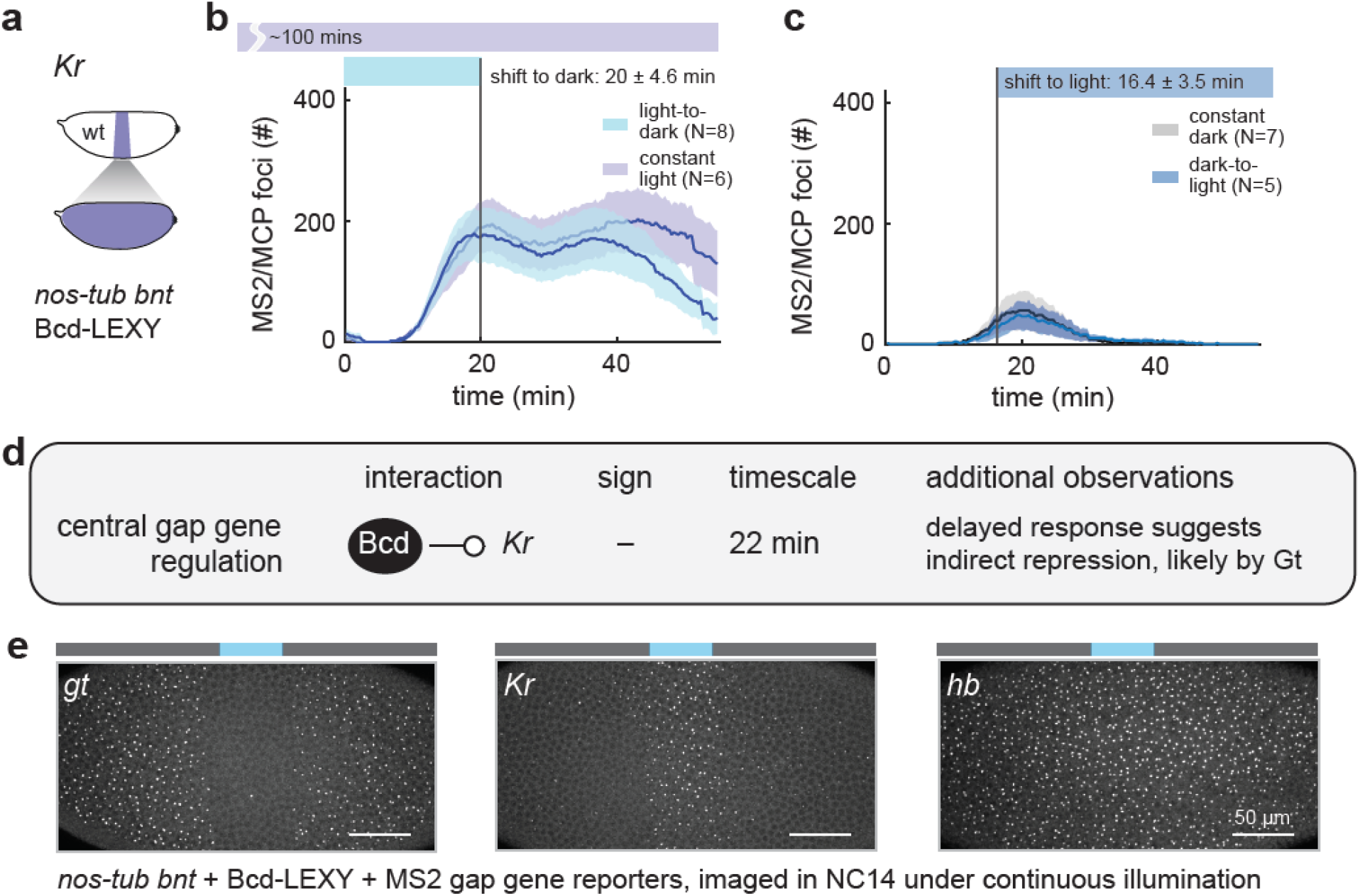
Acute perturbation of Bcd reveals delayed negative regulation of *Kr* transcription. (**a-c**) Optogenetic interrogation of Bcd-induced *Kr* transcription dynamics. Uniformly expressed Bcd-LEXY embryos were imaged using a *Kr* MS2 reporter (schematic in **a**) upon an acute shift from light to dark (in **b**) and dark to light (in **c**); constant-light and constant-dark stimuli were used as controls. (**d**) Summary of rapid perturbation results. An acute increase in nuclear Bcd-LEXY concentration drives loss of Kr expression after a 22 min delay. Conversely, *Kr* transcription is not observed for at least 1 h after an acute decrease in nuclear Bcd levels. (**e**) Measurement of *gt, Kr*, and *hb* transcription after continuous, local illumination in Bcd-LEXY embryos. Transcription of *gt* is suppressed where *Kr* is transcribed, whereas *hb* is largely unaffected in the Bcd-LEXY background. For **b-c** shaded regions show standard error of the mean, and the number of embryos tested is indicated on each plot.

We performed stimulus-response measurements in Bcd-LEXY embryos, which exhibit stark differences in *Kr* expression between constant blue light and dark conditions that reflect optogenetic switching across the anterior boundary of the *Kr* pattern (**Figure S4c**). An acute increase in nuclear Bcd-LEXY drove a corresponding decrease in *Kr* transcription, consistent with our expectation of low *Kr* expression at anterior positions (**Figure 5b**). However, unlike *hb* and *gt*, the change in *Kr* transcription only began after a 22 ± 2 min delay (**Table S2**).

Conversely, shifting the embryo from a high-Bcd state to a low-Bcd state in early NC14 did not lead to any detectable change in *Kr* expression prior to gastrulation, indicating that an even longer time period may be required to establish *Kr* expression upon loss of Bcd (**Figure 5c**).

These results support a model where high nuclear Bcd levels induce expression of a stable repressor of *Kr* expression (**Figure 5d**). An acute rise in Bcd would only produce the repressor after the time needed for new protein synthesis, and repressor degradation would be required for *Kr* expression to respond to a drop in Bcd activity. Our data also points to a likely candidate repressor among the gap genes. We observe a tight correlation between the spatial expression domains of *Kr* and *gt* in illuminated Bcd-LEXY embryos, suggesting that Gt may act as the long-lived, Bcd-induced negative regulator (**Figure 5e**). This model is well supported by prior studies^34-37^ identifying Gt as a potent repressor of *Kr* expression. In sum, our stimulus-response framework can be used to measure transcription dynamics that can in turn provide insight into direct and indirect links within a gene regulatory network.

### *kni* is transcribed rapidly upon light-triggered loss of nuclear Bcd

Our final target for optogenetic stimulus-response analysis was the posterior pattern of *kni* expression (**Figure 6a**). The gap gene *kni*, which is required for specification of posterior body segments, is thought to be induced by Bcd and Caudal (Cad) and repressed by Hb^38,39^. This complex and redundant regulation involves both maternally supplied anterior inputs (e.g., Bcd-dependent Cad patterning) and posterior cues (e.g., Nanos-dependent patterning of maternal Hb). Interestingly, we observe high *kni* expression even in *nos-tub bnt* embryos (**Figure 3b**), raising the question of how Bcd affects expression of a gap gene that is still highly expressed in the absence of Bcd.

**Figure 6.**
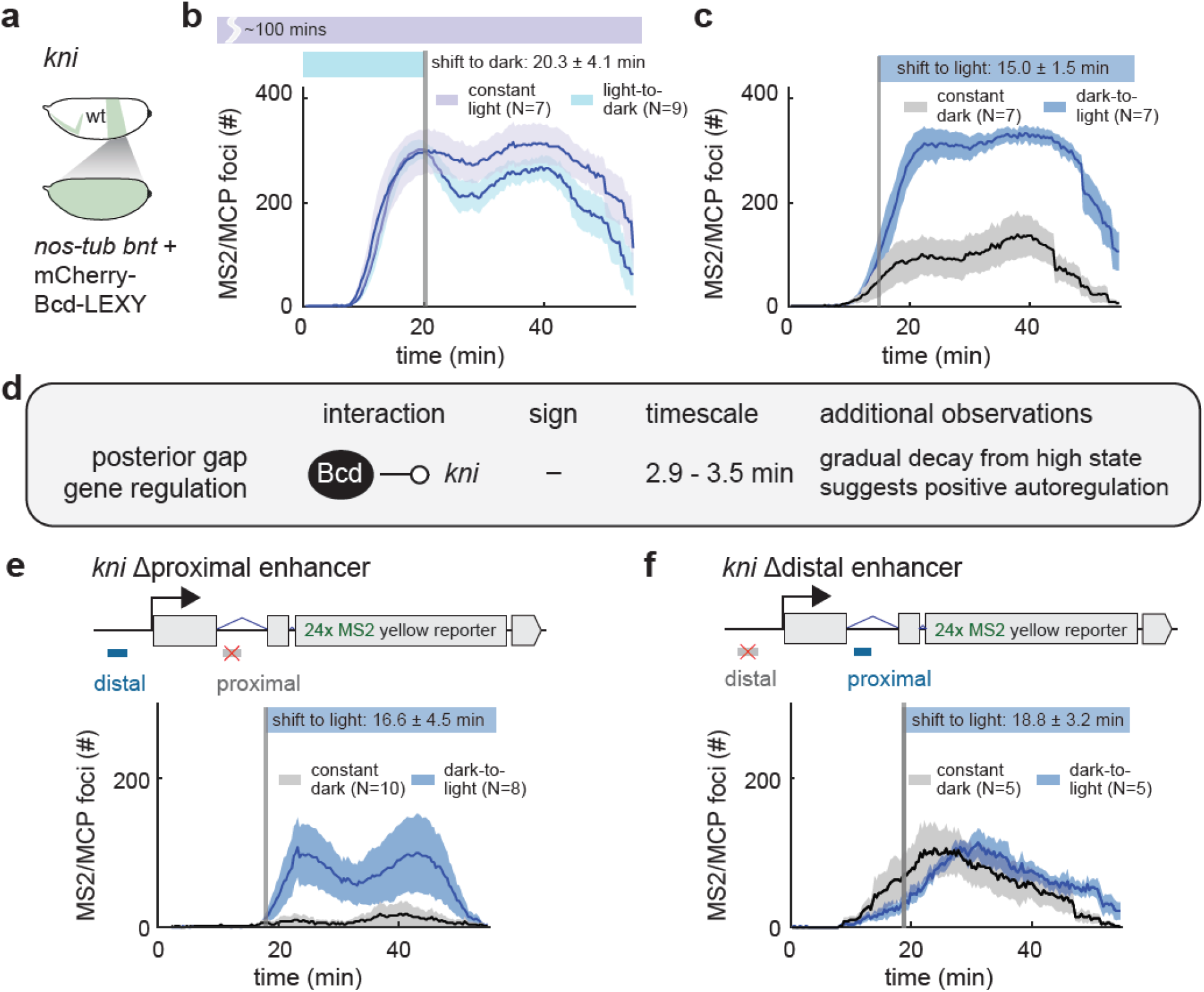
Acute removal of Bcd drives rapid activation of posterior *kni* transcription. (**a-c**) Uniformly expressed mCherry-Bcd-LEXY embryos were imaged using a *kni* MS2 transcription reporter upon an acute shift from light to dark (in **b**) and dark to light (in **c**); constant-light and constant-dark stimuli were used as controls. (**d**) Summary of rapid perturbation results. An acute decrease in mCherry-Bcd-LEXY expression, representing a change from central to posterior Bcd levels, drives a rapid rise in *kni* expression. In the converse experiment, *kni* transcription drops rapidly but only slightly upon acute Bcd nuclear import, suggesting that *kni* transcription is positively autoregulated. (**e**-**f**) Experiments as in **c** for *kni* reporters in which the proximal enhancer (in **e**) or the distal enhancer (in **f**) were replaced with non-targeted sequence. For **b–f**, shaded regions show standard error of the mean, and the number of embryos tested is indicated on each plot.

We examined the Bcd-dependent dynamics of *kni* expression^25^ in embryos expressing the lowest activity mCherry-Bcd-LEXY variant (**Figure 6b-c**; **Figure S4d**). Acutely dropping nuclear Bcd concentration led to a dramatic and unexpected change in *kni* transcription (**Figure 6c**). Within 2.9 ± 0.9 mins after a loss of nuclear Bcd, *kni* transcription began rising rapidly to levels that were comparable to those achieved under continuous illumination (**Figure 6b**). Conversely, a light-induced increase in nuclear Bcd triggered a similarly rapid but smaller-amplitude decrease in *kni* transcription (**Figure 6b**). Just as in the case of *gt*, the stability of the high-*kni*-expressing state may be indicative of positive autoregulation of *kni* expression by its own protein product. Together, these data suggest that Bcd can act as an apparent repressor of *kni* expression, an unexpected role for Bcd which is typically considered to perform only transcriptional activation functions. The initiation of *kni* transcription within 2 min after Bcd nuclear export is only compatible with a direct regulatory link, not Bcd-induced expression of an intermediate repressor.

To gain further insight into the repressive effect, we set out to define its requirements in the *kni* enhancer regions. The posterior pattern of *kni* expression is known to be regulated by two enhancers, an 818 bp proximal enhancer and a 2.3 kb distal enhancer^38,40^. We generated embryos expressing *kni* MS2 reporters with either the proximal or distal enhancer sequence replaced with non-regulated sequence^41^ and monitored expression in response to acute Bcd removal (**Figure 6e-f**). We found that the *kni* reporter lacking the proximal enhancer (*kni* Δproximal reporter) still showed potent regulation by mCherry-Bcd-LEXY, whereas the *kni* Δdistal reporter was not affected by light-induced changes in nuclear Bcd (**Figure S4e-f**; **Video S7-8**). Transcription from the *kni* Δproximal reporter also rose rapidly upon the shift to blue light, matching what was observed from the wild-type regulatory sequence (**Figure 6c,e**). Our results are consistent with prior observations that the *kni-*distal enhancer exhibits higher Bcd binding than does the *kni*-posterior enhancer, arguing that Bcd exerts its regulatory effects at the distal enhancer^38,40^. In summary, our acute stimulus-response framework identifies a rapid, repressive role for Bcd in regulating *kni* transcription through the *kni* distal enhancer, underscoring the power of optogenetic perturbation in a simplified genetic context to identify both known and unknown gene regulatory relationships.

## Discussion

### A stimulus-response strategy for dissecting complex developmental gene networks

We have described a combined genetic and optogenetic strategy to gain insight into one canonical developmental patterning network: control of gap gene expression by the Bicoid morphogen during *Drosophila* embryogenesis. Our strategy relies on three advances. First, we experimentally simplify the conditions under which the gap gene network operates, eliminating all pre-existing landmarks along the anterior-posterior axis to produce embryos with uniform positional identity. Although the reduced network involves just one input transcription factor (Bicoid) and four output genes (anterior *giant*, anterior *hunchback*, central *Krüppel*, and posterior *knirps*), it captures much of the complexity of the wildtype pattern including stripes of gap gene expression when Bcd is delivered in a head-to-tail gradient^2^. Second, we re-introduce optogenetic Bcd variants to shift these uniform embryos to any of three distinct anterior-posterior positions, enabling us to experimentally isolate specific gap genes patterns. Finally, we combine acute optogenetic perturbation with live-cell biosensors of target gene expression to map the network’s responses to acute changes in transcription factor concentration over time. Doing so required establishing new imaging methods for two-color confocal imaging and optogenetic activation *in vivo*, a challenge we solved by combining 970 nm two-photon imaging of GFP/mCherry with 450 nm excitation of the LEXY optogenetic system.

### Bcd-dependent regulation of anterior and posterior gap gene patterns

Our optogenetic stimulus-response experiments broadly support the canonical view of Bicoid as a direct transcriptional activator of *gt, hb*, and *Kr*. We find that both *gt* and *hb* are transcribed rapidly upon acute Bicoid nuclear import (**Figure 4**) and *Kr* expression requires the presence of low Bicoid activity (**Figure 5**). Our data also point to multiple regulatory links between gap genes. We find that *gt* expression is sustained when nuclear Bicoid is acutely removed (**Figure 4**), consistent with reports of its positive autoregulation^42^; that *Kr* exhibits delayed negative regulation by Bicoid (**Figure 5**), likely through Gt as an intermediate node^43^; and that *kni* expression remains high even after Bicoid nuclear import (**Figure 6**), consistent with its positive autoregulation^44^. Importantly, each of these network connections can be identified using a single, unified experimental workflow: acute optogenetic Bicoid perturbation and live recording and quantification of a target gene’s transcriptional dynamics.

Our study also revealed an unexpected result: rapid initiation of *kni* transcription after acute removal of mCherry-Bicoid-LEXY from the nucleus. Bicoid is not expected to act as a transcriptional repressor, so it is surprising to find any context in which its removal triggers rapid initiation of transcription. Classical models interpret loss of *kni* at anterior positions as being the consequence of indirect Bicoid-dependent regulation: repression by anterior gap gene products (e.g. *hb* or *Kr*) or weak activation by Caudal, which is translationally repressed by Bicoid^38,45,46^. Our result appears inconsistent with all of these explanations, as *kni* expression rises near-instantaneously after mCherry-Bicoid-LEXY nuclear export (**Figure 6**), too rapidly for changes in gap gene or Caudal protein levels to occur. Furthermore, rapid *kni* de-repression requires the distal enhancer, the predominant site of Bcd binding^38,40^.

How might rapid transcriptional activation occur upon loss of nuclear Bicoid? Our data is consistent with many possible mechanisms. Bicoid may compete for binding to the *kni* distal enhancer with another more potent transcriptional activator, such that Bicoid loss paradoxically increases *kni* expression. Alternatively, Bicoid may cooperatively associate with a transcriptional repressor at the *kni* enhancer and lead to increased repressor binding, mirroring the well-established interaction between Dorsal and Groucho for repressing subsets of genes along the dorsoventral axis^47,48^. We look forward to future studies that precisely define Bcd’s repressive role in *kni* expression, as well as extension of our acute stimulus-response methods to other complex gene regulatory networks.

Optogenetic stimuli have recently found widespread use in developmental contexts, from identifying critical time windows for developmental decisions^13,49-52^ to erasing and replacing signaling gradients with spatial light patterns^53^. However, optogenetic control on its own is not a panacea for revealing the inner workings of complex biological networks. Here we show that optogenetics can be used at a more granular level to home in on dynamic relationships between a transcription factor and its target genes *in vivo*. Nevertheless, work on an experimentally reduced system only constitutes a first step in understanding the full gap gene network, and we look forward to future studies that examine Bicoid-dependent responses as other factors from the natural system are systematically re-introduced. We can also envision extending the current approach to perturbing multiple nodes (e.g., by constructing LEXY fusions of all gap genes); quantitative modeling^4^ could elaborate network architecture still further. The future is bright for optogenetic interrogation of developmental gene networks.

## Material and methods

### Plasmids

Constructs were generated using In-Fusion assembly (Clontech) and oligonucleotides for primers were obtained from Integrated DNA Technologies. Constructs are available via Addgene or on request. Bcd-LEXY constructs are generated from pCol-*αTub67C-*EGFP-Bcd*-*FRT*-bcd 3’UTR-* 3xP3-RFP*-*FRT*-sqh 3’UTR*^54^, where the N-terminal EGFP was either removed or replaced by iRFP or mCherry and LEXY domain was inserted as C-terminus with a 15 amino-acid long linker in between. NLS-mCherry-LEXY constructs are generated by ligation of the NLS-mCherry-LEXY insert part PCR amplified from a mammalian expression vector (Addgene #72655^11^) and a fly expression vector pBabr-mTub-MCS-sqh3’UTR (courtesy from Wieschaus lab) digested by restriction enzymes NheI and SpeI. mCherry was subsequently replaced by EGFP to generate NLS-EGFP-LEXY plasmid.

### Fly stocks and genetics

#### Establishing Bcd-LEXY and *bcd nos tsl* fly stocks

For generation of transgenic flies and stocks, all four Bcd-LEXY (Bcd-LEXY, EGFP-Bcd-LEXY, iRFP-Bcd-LEXY and mCherry-Bcd-LEXY) constructs were integrated into the third chromosome using the ϕC31-based integration system^55^ at the VK33 site (65B2) by BestGene. NLS-mCherry-LEXY and NLS-EGFP-LEXY constructs were integrated into the second chromosome at the VK02 site (47C6). Each Bcd-LEXY variant was then further recombined either with *bcd*^*E1*^ or *bcd*^*E1*^ *nos*^*BN*^ *tsl*^*4*^ on the third chromosome^54^. NLS-mCherry-LEXY was recombined with MCP-mNeonGreen on the second chromosome, and further crossed with *bcd*^*E1*^ *nos*^*l7*^ *tsl*^*4*^ on third chromosome to generate MCP-mNeonGreen NLS-mCherry / Cyo; *bnt* / TM3 flies.

We obtained few and poor-quality embryos from *bcd*^*E1*^ *nos*^*BN*^ *tsl*^*4*^ homozygous females, and thus used *bcd*^*E1*^ *nos*^*BN*^ *tsl*^*4*^ */ bcd*^*E1*^ *nos*^*l7*^ *tsl*^*4*^ transheterozygotes for further experiments. While *nos*^*BN*^ is a complete loss of both *nos* RNA and protein, which impedes both pole cell migration and therefore germline cells formation and abdominal segmentation, *nos*^*l7*^ is a partial deletion near C-terminal of the zinc-finger domain that maintains normal germline development. However, in terms of body segmentation phenotype and gap gene expression pattern, *nos*^*BN*^ and *nos*^*l7*^ both exhibit indistinguishable patterns as expected from severe loss of function^56,57^, supporting the use of this transheterozygous background for *nanos* loss of function in A-P patterning.

#### Establishing uniform Bcd-LEXY embryos

To achieve uniform Bcd expression, Bcd-LEXY, *bnt* flies were crossed to heat shock-inducible *flippase* expressing flies and the resulting larvae were heat shocked at 37°C for three continuous days for 1 hour each day. After one generation of outcrossing, progeny lacking the *bcd* 3’UTR were sorted by loss of RFP expression. Then Bcd-LEXY variants were driven by *sqh* 3’UTR (see **Figure 2A**) resulting in a uniform distribution of Bcd along the AP axis.

#### Establishing *nos-tub bnt* uniform Bcd-LEXY embryos

In wild-type embryos, *nanos* mRNA is localized at the posterior pole and produces a posterior-to-anterior gradient of Nanos protein^19^. A second population of *nanos* mRNA is not asymmetrically patterned and produces uniform Nanos protein that plays a crucial role in suppressing maternal Hunchback translation^20^. Complete loss of *nanos* disrupts both the patterned and uniform contributions, leading to abnormally high levels of Hunchback throughout the embryo. We thus used a nos-tub:TCEIIUC:AG construct^20^ (courtesy of the Gavis lab) as a uniformly-expressed, reduced-activity form of Nanos to reduce maternal Hb levels, thereby allowing expression of abdominal gap genes like *kni* and *gt*. The construct was further recombined to the Sp marker on the same chromosome to mark the transgene, and then crossed with male uniform Bcd-LEXY *bcd*^*E1*^ *nos*^*BN*^ *tsl*^*4*^ / TM3 flies to generate nos-tub:TCEIIUC:AG /+; uBcd-LEXY *bcd*^*E1*^ *nos*^*BN*^ *tsl*^*4*^ / TM3 flies. By crossing males of the preceding genotype to MCP-mNeonGreen, NLS-mCherry / Cyo; *bcd*^*E1*^ *nos*^*l7*^ *tsl*^*4*^ / TM3 females, we obtained female flies with the genotype nos-tub:TCEIIUC:AG / MCP-mNeonGreen NLS-mCherry; uBcd-LEXY *bcd*^*E1*^ *nos*^*BN*^ *tsl*^*4*^ / *bcd*^*E1*^ *nos*^*l7*^ *tsl*^*4*^ that we then caged with MS2 reporter males for live imaging.

### Cuticle preparation

For dark and light conditions, embryos with specific Bcd-LEXY variants were collected between 0-1 h post laying in the dark on an agar plate. Then embryos for the light condition were placed under a custom-built panel of blue LEDs and removed from light after 4hrs. In dark conditions, embryos were kept in a light-tight box away from ambient room light or blue light to prevent inadvertent optogenetic stimulation. After a 3 h incubation in light or dark conditions, embryos were kept at room temperature (at 22°C) for another 24-36 h and then bleached, then moved to the methanol-heptane glass tube and vigorously shaken for 20 sec. Embryos settled at the bottom were removed and placed on a glass slide with Hoyer’s solution (1:1 premix lactic acid) and sandwiched between the glass slide and cover glass. The slide was placed at 65°C overnight and then imaged on a Nikon Eclipse Ni dark-field microscope at 10x zoom.

### Immunostaining and imaging

Embryos were collected every 2 h and aged in dark for another 2 h. Embryos were dechorionated by bleaching and then heat fixed and stained essentially as described in^1,54^ with rabbit anti-Bcd, mouse anti-Hb primary antibodies (courtesy by Eric Wieschaus) and sheep anti-GFP (Invitrogen, USA) followed by fluorophore-conjugated secondary antibodies Alexa-488 (sheep), Alexa-555 (mouse), and Alexa-647 (rabbit) from Invitrogen. For pairwise comparisons of wild-type and mutant backgrounds, embryos expressing HisGFP collected the same way were mixed in each tube for staining and imaging. Stained embryos were imaged on a Nikon A1R laser-scanning confocal microscope, and a 5 μm z-stack around the midsagittal plane with step size of 1 μm were taken. For data analysis, custom MATLAB imaging analysis code recognized the contour of the embryo and extracted the intensity of the surface nuclei for all three channels. Intensities of three channels were normalized to HisGFP embryos that mixed in each slide respectively, with 1 being the mean maximal intensity of HisGFP embryos. For Hb level, min-max normalization was further conducted for clear comparison of boundary position, and half maximal positions of the posterior boundary of anterior expression domain were picked out for each genotype for the box plot.

### Two-photon microscopy

A custom microscope was built to simultaneously perform two-photon excitation imaging and localized optogenetic stimulation on the same setup. A Chameleon Ultra II tunable laser was used at 970 nm to simultaneously excite green and red-tagged biomolecules^17,58,59^. The laser beam was collimated and passed through a laser power modulator Pockels cell (350-80-LA-02 KD P Series E-O Modulator, Conoptics, USA). The output laser beam was expanded to 4 mm diameter (AC254-050-AB-ML, AC254-150-AB-ML, Thorlabs, USA) before reaching a two-axis scan galvo mirror (6210H, Cambridge Technology, USA). After the scan mirrors, the laser beam passed through a f-theta lens (focal length 63 mm; 4401-388-000-20, Linos, USA), a tube lens (focal length 180 mm, AC508-180-AB-ML, Thorlabs) and focused on the imaging sample using a high numerical aperture objective (NA, Nikon 1.3 NA, 40X). The fluorescence signal was collected and sent to two sensitive point photo multiplier tubes (H10770A-40, Hamamatsu, Japan). The microscope setup interfaces via data acquisition cards (DAQ; PCIe 6321 and PCIe 6374, National Instruments, USA) using MATLAB-based ScanImage 5.6 software^60^. For live imaging, embryos were imaged close to cover glass surface (image resolution 1024×512 pixels at 3.2 μs pixel dwell-time; see imaging details in **Table S3**).

### Optogenetic stimulation

LEXY perturbation was achieved using a digital micro-mirror device (DMD; DLP 4500 LightCrafter, Texas Instruments, USA) to project spatial patterns and to rapidly change light levels^5^ (see **Figure S2a**) through a parallel light path using a long-pass 473 nm dichroic mirror and a combination of color and interference filters to attenuate the DMD’s blue LED wavelength (445 ± 8 nm). To synchronize two-photon image acquisition and DMD blue light activation cycles, an external trigger mode in DLP LightCrafter control software was used. The software controls the LED light wavelength, pulse duration, pulse duty cycle, the number of pulses, and the type of spatial image pattern to project on the imaging sample (see details in **Table S3**). The optimum blue light level for optogenetic perturbation was determined by optimizing the maximum protein export with minimal light scattering to neighboring nuclei (see **Figure 2**e). After scanning the range between 50–250 μW/cm^2^ of blue-light on/off pulsatile cycles of LEXY-tagged protein nuclear signal (data not shown), 100 μW/cm^2^ was determined for all optogenetic perturbations performed in this study.

### Live imaging data collection

For the live data acquisition and light perturbation experiments, flies were kept in dark at 25°C and the embryos were collected on an agar plate between 1-2 hrs post laying. For live imaging, embryos were dechorionated on double-sided tape and mounted on a glued membrane film (Lumox film, Starstedt, Germany). Followed by covering them in halocarbon oil 27 and sandwiched between the membrane and the cover glass slide (cover glass washed and cleaned with pure ethanol, slide #1.5, Sigma BR470045). The data collection was performed using a custom-built two-photon microscope using 970 nm laser excitation for green (EGFP and mNeonGreen tagged protein and MS2 loops respectively) and red (only the NLS-mCherry-LEXY) at room temperature (ranges from 21.5-22.5 °C). And the blue light perturbation was done by a DMD unit installed on the same setup; see (see microscope section). Summarized details on data collection for specific experiments are tabulated in **Table S3**.

### Data quantification and statistical analysis

#### Bcd-LEXY activity

To estimate the functional Bicoid activity (potency) of fluorescently tagged Bcd-LEXY fusion proteins (as well as remaining activity in *bcd*^*E1*^ homozygous mutant fly lines and the Bcd dose level), we used protein immunostaining of the Bcd target gene Hb and measured position shifts in the posterior Hb boundary as well as position shifts of the cephalic furrow. All shifts were scaled according to embryo length and the quantified estimates are presented in **Figure 2d-e** (**Table S1** for cephalic furrow position shifts).

#### LEXY tagged protein export and import kinetics

Blue light-induced LEXY export kinetics and nuclear localization signal (NLS)-induced import kinetics were determined by analyzing the nuclear intensity of the fluorescent moiety of these fusion proteins. Intensity time traces were averaged and fitted with a single exponential n0*exp(-t/τ) to estimate the export rate, i.e. the inverse of the time constant. Similarly, the import time constants were estimated using n0*(1-exp(-t/τ)) as the fitting model (**Figure 2f-h**). Note: the uniform EGFP-Bcd-LEXY, NLS-mCherry-LEXY, and NLS-EGFP-LEXY lines were measured using the DMD-equipped custom-built two-photon microscope, while the uniform mCherry-Bcd-LEXY and iRFP-Bcd-LEXY lines were imaged on a commercial Nikon A1R confocal microscope (imaging conditions can be found in **Table S3**). For LEXY translocation kinetics fits, the mean and the standard deviation are presented for multiple embryo replicates.

#### Quantification of the reporter gene MS2 spots

We created a custom MATLAB script to analyze and visualize time-lapse MS2 counts. In brief, two-color raw tiff image data was acquired on a custom-built two-photon microscope (MS2 data acquisition setting details can be found **Table S3**; see **Figure S3** and **Figure S4**). The red channel NLS-mCherry-LEXY nuclear signal was segmented, and intensity traces were used to estimate nuclear cycle start and the optogenetic perturbation state during MS2 imaging. The rise in NLS-mCherry-LEXY signal time was set to 5.6 min in NC14 (this x-axis time offset was set for C14 start for both red and green channels; see **Figure S3d**). For MS2 data, the image data was z-max projected (8 total z slices, each 1.1 μm apart), then a 2D gaussian filter was applied to filter-out small structures and followed by threshold to select MS2 spots. Finally, every spot was counted as an active MS2 spot in the center part of the embryo (40×150 μm^2^ ROI) for either blue light illuminated or dark conditions embryos. All the data representing spot count time traces indicate the mean and standard error of the mean over multiple embryo measurements, unless stated otherwise.

#### Mean response time post light perturbation

For response time quantification, we measured the difference between individual MS2 foci count time traces (On-Off) with mean MS2 foci count time traces (On) light condition (**Figure 4b** light blue color individual traces subtracted by mean purple trace). Followed by taking gradient (25% amplitude change) and embryos showing no change or below the threshold (<25%) were excluded from the analysis. Finally, the response mean-time and standard error of the mean were reported in **Table S2** (taking account of error in light on/off time by using error propagation).

## Acknowledgements

The authors thank members of the Gregor and Toettcher laboratories, Liz Gavis, Mustafa Khammash, Sant Kumar, Jason Puchala, and Trudi Schüpbach. The project was supported by NSF grant PHY–1734030 (T.G.) and CAREER–1750663 (J.E.T.); NIH grants R01GM097275 (T.G.), U01DA047730 (T.G.), U01DK127429 (T.G. and J.E.T.), T32GM007388 (M.S.); and a Princeton MOL Innovation Award (J.E.T.). We also acknowledge imaging support from the Princeton Molecular Biology Microscopy Facility, which is a Nikon Center for Excellence.

