## Supplementary Figures and Tables for "Optogenetic control of the Bicoid morphogen reveals fast and slow modes of gap gene regulation"

**This PDF file includes:**


| | |
|---|---|
| Supplementary Figures 1-4 | Pages 2-5 |
| Supplementary Tables 1-3 | Pages 6-8 |
| Supplementary Video Legends | Page 9 |
| Supplementary References | Page 9 |

### Supplementary Figures

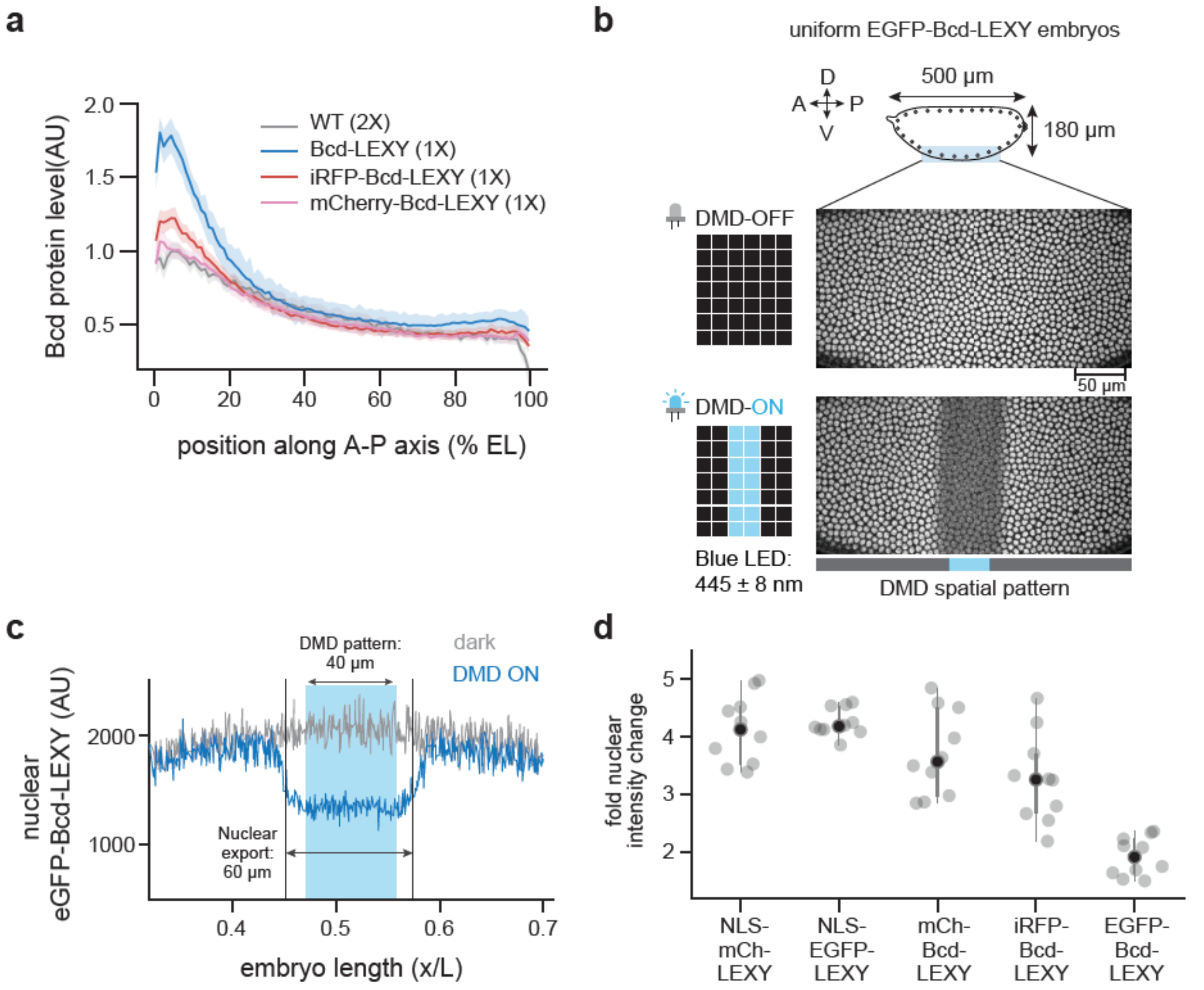


**Figure S1. Characterization of Bcd-LEXY expression and translocation spatial precision and dynamics in different variants.** (**a**) Immunofluorescence for Bcd protein for three Bcd-LEXY variants, compared to WT embryos. Bcd levels are quantified as a function of position and in the genetic background of a single copy of gradient αTub67C>Bcd-LEXY expressed in *bcd*$^{E1}$ homozygous embryos. (**b-c**) Spatial precision of EGFP-Bcd-LEXY import and export (**Video S3**). Representative images are shown in **b**, where the top panel shows the unilluminated state and bottom panel has a central stripe DMD pattern of 40 µm wide. As quantification of the pattern shown in **c**, the actual pattern on the embryo is 60µm wide, which is ~10-12µm wider (about one cell width) than the DMD pattern on both sides. (**d**) Nuclear fluorescence fold changes between dark and light conditions for different LEXY variants. Fold change is quantified by division of background subtracted nuclear fluorescent intensity in dark condition by light condition. *Related to Figure 2.*

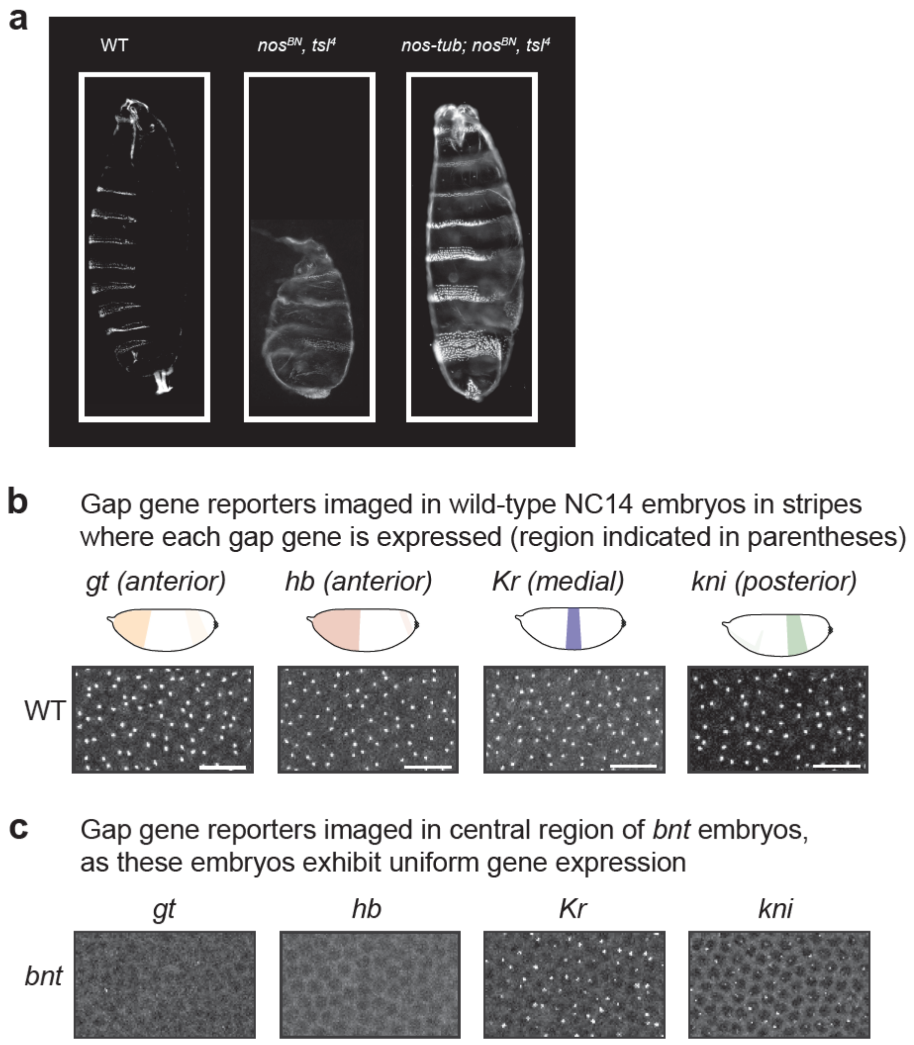


**Figure S2. Phenotypic and genotypic characterization in WT, *nt* and *bnt* embryos.** (**a**) Larval cuticles of $nos^{BN}$*, tsl*$^{4}$ double mutant embryos and $nos^{BN}$*, tsl*$^{4}$ double mutants with low uniform Nanos expression(*nos-tub*). $nos^{BN}$*, tsl*$^{4}$ double mutant larva formed only 1-2 abdominal segments due to high Hb level at posterior part, whereas a low uniform expression of Nanos restores abdominal segments by repressing Hb translationally. (**b**) Characterization of gap gene expression in WT and *bnt* embryos via MCP/MS2 imaging. Images shown are 43.9µm × 73.5µm representative regions where four gap genes are expressed correspondingly, captured around 25-35min into NC14. Compared to *nos-tub bnt* embryos (**Figure 3b**), *bnt* embryo has low *kni* expression and barely no *gt* expression, but higher *Kr* expression, representing a more anterior embryonic position due to the presence of maternal *hb*. *Related to Figure 3.*

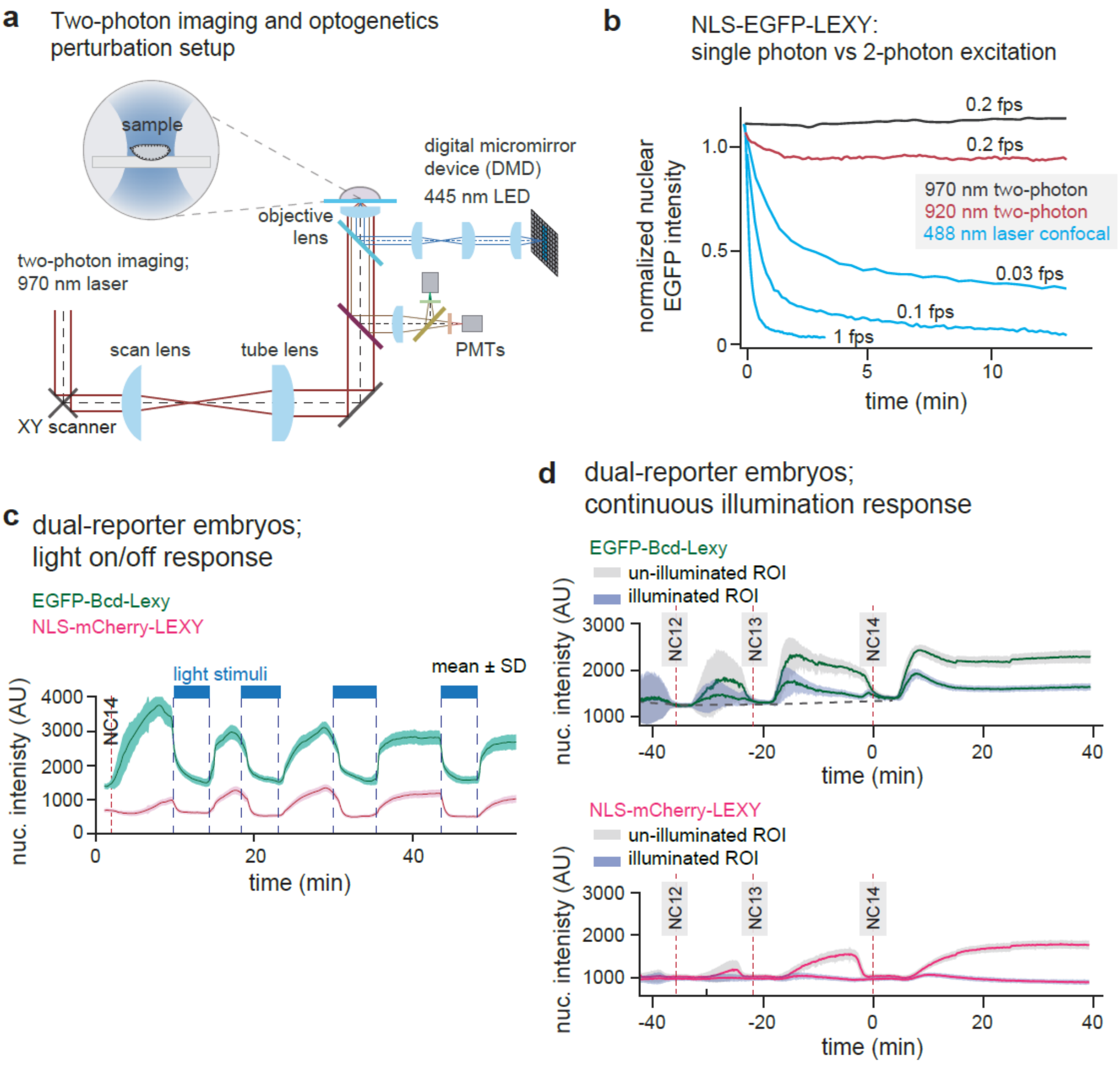


**Figure S3. Two-photon setup for dual-color high-resolution imaging and optogenetic perturbation.** (**a**) Microscope schematic of a two-photon laser confocal for live imaging and the DMD unit for optogenetics perturbations. (**b**) Comparison of single-photon 488 nm laser excitation and two-photon imaging of NLS-EGFP-LEXY. Whereas single-photon 488 nm laser and two-photon 920 nm laser still trigger activation of Aslov2 domain, thus resulting in nuclear export of NLS-EGFP-LEXY, two-photon 970 nm laser has no detectable effect on NLS-EGFP-LEXY. (**c**) Fast and reversible control of NLS-mCherry-LEXY and EGFP-Bcd-LEXY import and export in pulsatile cycles in NC14. Both constructs were co-expressed in the same embryo. (**d**) Quantification of EGFP-Bcd-LEXY and NLS-mCherry-LEXY during NC11-14 in constant dark and light conditions. In dark condition, NLS-mCherry-LEXY enters nuclei gradually after each division in contrast to EGFP-Bcd-LEXY.Furthermore, even in continuous blue light exposure, EGFP-Bcd-LEXY enriches in nuclei transiently right after division. This may be due to that Bcd tends to stick to DNA and right after nuclear envelope reforms, Bcd-LEXY might be brought into nuclei with DNA, which would potentially be faster than regular nuclear import mediated by NLS. For **b-c** shaded regions show standard error of the mean across nuclei in the same embryo. *Related to Figure 4.*

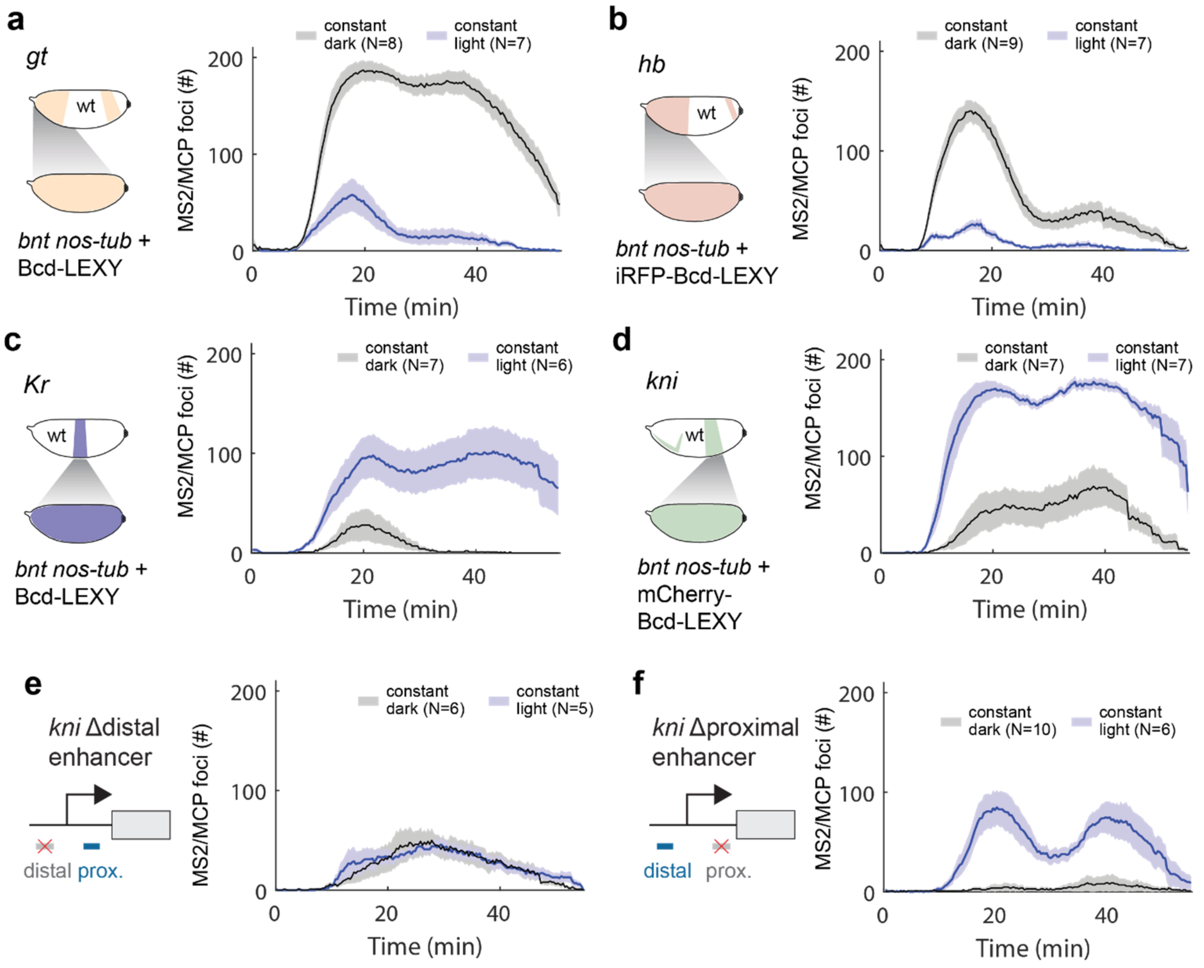


**Figure S4. Live transcription measurement of gap genes in constant dark and light conditions.** (**a-b**) Quantification of *hb* and gt transcriptional activity in constant dark and light conditions via MCP/MS2.*hb* and *gt* show high expression in dark and low expression in constant blue light, indicating that Bcd acts as a positive regulator for *hb* and *gt* (**Video S3, S4)**. (**c-d**) Quantification of *hb* and *gt* transcriptional activity in constant dark and light conditions via MCP/MS2(**Video S5, S6)**. In contrast to *hb* and *gt*, *Kr* and *kni* MS2 show higher expression constant light than dark condition, indicating that Bcd may act as a negative regular for *Kr* and *gt*. (**e-f**) *kni* Δproximal and *kni* Δdistal enhancer driven MS2 in constant dark and light conditions. *kni* Δproximal enhancer has much higher transcription activity in light compared to dark, similar to full *kni* BAC MS2 in **d**, whereas *kni* Δdistal enhancer barely has any changes. Thus, *kni* distal enhancer is mainly responsible for the change observed via optogenetic control. For **a-f** shaded regions show standard error of the mean, and the number of embryos tested is indicated in the legend of each plot. *Related to Figures 4-6*.

#### Supplementary Tables

**Table S1. Hatch rates and cephalic furrow positions in gradient optogenetic Bcd-LEXY variants.**

| Genotype | Hatch rate, dark | Hatch rate, light | Cephalic furrow position, dark<br>Mean ± SD(%EL) | Cephalic furrow position, light<br>Mean ± SD(%EL) |
|---|---|---|---|---|
| WT | 92.3% (192/209) | 87.9% (182/207) | 33.2±1.7 (N=8) | 33.6±1.4 (N=10) |
| Bcd-LEXY, $bcd^{E1}/bcd^{E1}$ | 69.9% (269/385) | 0% (0/49) | 37.5±1.4 (N=12) | 26.1±1.9 (N=11) |
| iRFP-Bcd-LEXY, $bcd^{E1}/bcd^{E1}$ | 42.0% (131/312) | 0% (0/25) | 27.1±2.4 (N=16) | Not formed |
| mCherry-Bcd-LEXY, $bcd^{E1}/bcd^{E1}$ | 0% (0/78) | 0% (0/22) | Not formed | Not formed |

**Table S2. Estimates of mean response time post light perturbation**

| **MS2 reporter** | **Mean response time post light perturbation mean ± SEM** | |
|---|---|---|
| | **Light On- Off transition** | **Light Off- On** |
| *hb* | 1.7 ± 0.9 (11/12) ; **Figure 4d** | 3.3 ± 1.1 (9/10); ; **Figure 4e** |
| *gt* | 5.9 ± 1.6 (6/9); **Figure 4g** | 9 ± 2.7 (5/6) **Figure 4h** |
| *Kr* | 21.7 ± 1.8 (7/8); **Figure 5b** | NA (no change); **Figure 5c** |
| *kni* | 3.5 ± 1.7 (7/9); **Figure 6b** | 2.9 ± 0.9 (7/7); **Figure 6c** |

#### Table S3. Live protein and mRNA reporter lines and imaging parameters

| Reporter line | Imaging conditions | DMD light conditions |
|---|---|---|
| αTub67C> NLS-EGFP-LEXY (**Figure 2h**) | Custom built two-photon microscope<br>laser-970 nm<br>laser power- 10 mW before objective<br>z-slice step= 1.1 μm<br>total z-slices= 2<br>single time frame=5 sec<br>pixel size=300 nm Image resolution =1024x512 pixels | DMD blue light wavelength= 445 nm<br>Pulse duration=40 ms<br>Pulse duty cycle= 100 ms<br>Number of pulses= 5<br>Blue light power 100 $\mu$W/cm$^2$ |
| αTub67C> NLS-mCherry-LEXY (**Figure 2h**) | " | " |
| αTub67C> EGFP-uBcd-LEXY (**Figure 2h**) | " | " |
| αTub67C> iRFP-uBcd-LEXY (**Figure 2h**) | **Nikon A1R** laser-scanning confocal<br>640 nm 20X air objective<br>laser power - 10% ,<br>PMT HV 120, PMT offset 8<br>time interval = 5 sec<br>pixel size=300 nm<br>image resolution =1024x512 pixels | DMD blue light wavelength= 445 nm |
| αTub67C> mCherry-uBcd-LEXY (**Figure 2h**) | 561 nm<br>rest same as above | " |

| | | |
|---|---|---|
| *gt* (**Figure 4**) | custom built two-photon microscope<br>laser-970 nm<br>laser power- 10 mW before objective<br>z-slice step= 1.1 μm<br>total z-slices= 8<br>single time frame=20 sec<br>pixel size=300 nm<br>image resolution = 1024x512 pixels | DMD blue light wavelength= 445 nm<br>Pulse duration=40 ms<br>Pulse duty cycle= 100 ms<br>Number of pulses= 5<br>Blue light power 100 $\mu$W/cm$^2$ |
| *hb* BAC (**Figure 4**) | " | " |
| *Kr* (**Figure 5**) | " | " |
| *kni* BAC (**Figure 6**) | " | " |
| *kni* BAC ΔDist (**Figure 6**) | " | " |
| *kni* BAC ΔProx (**Figure 6**) | " | " |

#### Table S4. Key resources

| Fly stocks | Source | Reference/Details |
|---|---|---|
| $bcd^{E1}$ $nos^{BN}$ $tsl^{4}$ | Wieschaus/Schüpbach | Princeton University fly stock |
| $bcd^{E1}$ $nos^{L7}$ $tsl^{4}$ | Wieschaus/Schüpbach | Princeton University fly stock |
| Sp, nos-tub | Wieschaus | Princeton University fly stock |
| nos>ϕ NLS-MCP-mNeonGreen | This study | |
| αTub67C> NLS-mCherry-LEXY | This study | |
| αTub67C> NLS-eGFP-LEXY | This study | |
| αTub67C> eGFP-Bcd-LEXY FRT bcd 3' UTR hsp70 RFP FRT sqh 3'UTR | This study | |
| αTub67C> Bcd-LEXY FRT bcd 3' UTR hsp70 RFP FRT sqh 3'UTR | This study | |
| αTub67C> iRFP-Bcd-LEXY FRT bcd 3' UTR hsp70 RFP FRT sqh 3'UTR | This study | |
| αTub67C> mCherry-Bcd-LEXY FRT bcd 3' UTR hsp70 RFP FRT sqh 3'UTR | This study | |
| hb- MS2 BAC | NA | (Bothma et al., 2015) |
| Kr- MS2 reporter | NA | (El-Sherif and Levine, 2016) |
| kni- MS2 BAC | NA | (Bothma *et al.*, 2015) |
| kni ΔDistal- MS2 BAC | NA | (Bothma *et al.*, 2015) |
| kni ΔProx- MS2 BAC | NA | (Bothma *et al.*, 2015) |
| gt- MS2 reporter | NA | (Syed et al., 2021) |
| **Software and control unit** | | |
| DLP 4500 LightCrafter control software | DMD control software | Texas Instruments |
| DLP 4500 LightCrafter unit | Image projection device | Texas Instruments |
| Matlab | Image and data processing | Mathworks |
| Fiji | Image processing | Fiji |
| National Instruments/Labview | Imaging control synchronization | National Instruments |

#### Supplementary Video Legends

**Video S1-S4 Blue light induced differential gap gene expression by controlling Bcd concentration.** Maximum intensity projection time-lapse movie of embryos expressing a single uniformly-expressed Bcd-LEXY variant in the *nos-tub bnt* genetic background with a single gap gene MS2/MCP mRNA biosensor. Each video shows transcriptional responses for a different gap gene and Bcd-LEXY variant: *hb* in iRFP-Bcd-LEXY (**Video S1**); *gt* in Bcd-LEXY (**Video S2**); *Kr* in Bcd-LEXY (**Video S3**); and *kni* in mCherry-Bcd-LEXY (**Video S4**). A stripe of blue light was projected on the center part of the embryo, with light treatment initiated in NC10-NC11 and maintained until the end of NC14. See live imaging data acquisition parameters in **Table S3**. *Related to Figures 3-6.*

**Video S5 EGFP-Bcd-LEXY imaged using 970nm two-photon excitation without blue light perturbation.** A representative embryo expressing uniform EGFP-Bcd-LEXY, $bcd^{E1}/bcd^{E1}$ imaged during nuclear cycle 14 on a custom built two-photon microscope at 970 nm. Single plane time-lapse images were acquired every 5 sec close to the glass surface without any blue light treatment. The movie shows a stable nuclear localization signal of Bcd and no loss in signal was observed during the imaging process(see **Figure S2b**). The video frame visualization runs at 40 fps. *Related to Figure 4.*

**Video S6 Spatial protein EGFP-Bcd-LEXY pattern created by using blue light.** Continuous blue light induced Bcd protein spatial pattern shaped by exporting EGFP-Bcd-LEXY out of the cell nucleus (genotype EGFP-Bcd-LEXY, $bcd^{E1}/bcd^{E1}$; see **Figure S2c, d** for spatial quantification and **Figure S3b-top** for temporal quantification). The images were acquired every 5 sec and the video frame visualized at 40 fps. *Related to Figure 4.*

**Video S7-8. Probing enhancer control of Bcd-dependent *kni* repression using reporters lacking proximal and distal enhancer elements.** Maximum intensity projection time-lapse movie of embryos uniformly expressing mCherry-Bcd-LEXY in the *nos-tub bnt* genetic background. Embryo also express an MS2/MCP biosensor containing the *kni* regulatory region without its proximal enhancer (Δproximal; **Video S7**) or without its distal enhancer (*kni*Δdistal; **Video S8**). A stripe of blue light was projected on the center part of the embryo, with light treatment initiated in NC10-NC11 and maintained until the end of NC14. See live imaging data acquisition parameters in **Table S3**. *Related to Figure 6.*
